# Comparative infectivity and pathogenesis of emerging SARS-CoV-2 variants in Syrian hamsters

**DOI:** 10.1101/2021.02.26.433062

**Authors:** Rana Abdelnabi, Robbert Boudewijns, Caroline S. Foo, Laura Seldeslachts, Lorena Sanchez-Felipe, Xin Zhang, Leen Delang, Piet Maes, Suzanne J. F. Kaptein, Birgit Weynand, Greetje Vande Velde, Johan Neyts, Kai Dallmeier

## Abstract

Within one year after its emergence, more than 108 million people contracted SARS-CoV-2 and almost 2.4 million succumbed to COVID-19. New SARS-CoV-2 variants of concern (VoC) are emerging all over the world, with the threat of being more readily transmitted, being more virulent, or escaping naturally acquired and vaccine-induced immunity. At least three major prototypic VoC have been identified, i.e. the UK (B.1.1.7), South African (B.1.351) and Brazilian (B.1.1.28.1), variants. These are replacing formerly dominant strains and sparking new COVID-19 epidemics and new spikes in excess mortality. We studied the effect of infection with prototypic VoC from both B.1.1.7 and B.1.351 lineages in Syrian golden hamsters to assess their relative infectivity and pathogenicity in direct comparison to two basal SARS-CoV-2 strains isolated in early 2020. A very efficient infection of the lower respiratory tract of hamsters by these VoC is observed. In line with clinical evidence from patients infected with these VoC, no major differences in disease outcome were observed as compared to the original strains as was quantified by (*i*) histological scoring, (*ii*) micro-computed tomography, and (*iii*) analysis of the expression profiles of selected antiviral and pro-inflammatory cytokine genes. Noteworthy however, in hamsters infected with VoC B.1.1.7, a particularly strong elevation of proinflammatory cytokines was detected. Overall, we established relevant preclinical infection models that will be pivotal to assess the efficacy of current and future vaccine(s) (candidates) as well as therapeutics (small molecules and antibodies) against two important SARS-CoV-2 VoC.

## Introduction

Barely one year after surfacing and global spread of SARS-CoV-2, more than 113 Mio infected people and 2.5 Mio fatal cases have been reported worldwide (as of February 26^th^ 2021). Variants of SARS-CoV-2 variants are emerging in different parts of the world, posing a new threat. Currently at least three major prototypic virus variants of concern (VoC) have been detected respectively in the UK (lineage B.1.1.7;^1–3^ earliest sample date 2020-02-03), South Africa (B.1.351 or 501Y.V2;^1,4,5^ earliest sample date 2020-10-08) and Brazil (B.1.1.28.1 or P.1;^6,7^ earliest sample date 2020-12-15). Even in highly endemic regions, these VoC started to replace formerly dominant strains and appear to be at the root of new waves of infections and new spikes in excess mortality, raising the concern of being more readily transmitted or of being even more virulent. In general, numerous mutations have been identified in the VoC genomes, which occur in different coronavirus genes. Though, the Spike (S) protein appears to be particularly prone to accumulate amino acid changes. As such the VoC of UK, South African and Brazil origin share the prominent mutation N501Y in the receptor-binding domain (RBD) of S that may favor viral attachment to its cellular receptor hACE2.^8^ Moreover, massive outbreaks in previously heavily affected regions, such as in the Manaus area of Northern Brazil with seropositivity rates of up to 76% prior to disease resurgence, raise the concern of VoC escaping pre-existing immunity.^7^ Likewise, several vaccine candidates either failed to show efficacy, or at least displayed a marked drop in vaccine efficacy in Phase 3 clinical trials in regions of South Africa where the VoC B.1.351 is circulating.^9,10^ Escape from antibody neutralization has been linked to amino acid changes in key residues of S, such as E484K, found in both the South African and Brazilian variant and as well in some more recently emerging UK (sub)lineages.^11–13^ The evolution of such new SARS-CoV-2 VoC may be driven by viral escape under host immune pressure during acute infection,^14^ and resulting SARS-CoV-2 antigenic drift is feared to spark future COVID-19 epidemics.

Here, we investigate infection of Syrian hamsters with prototypic VoC, namely local Belgian low-passage isolates from both B.1.1.7 and B.1.351 lineages (Fig 1A, B).^15–17^

**Fig 1.**
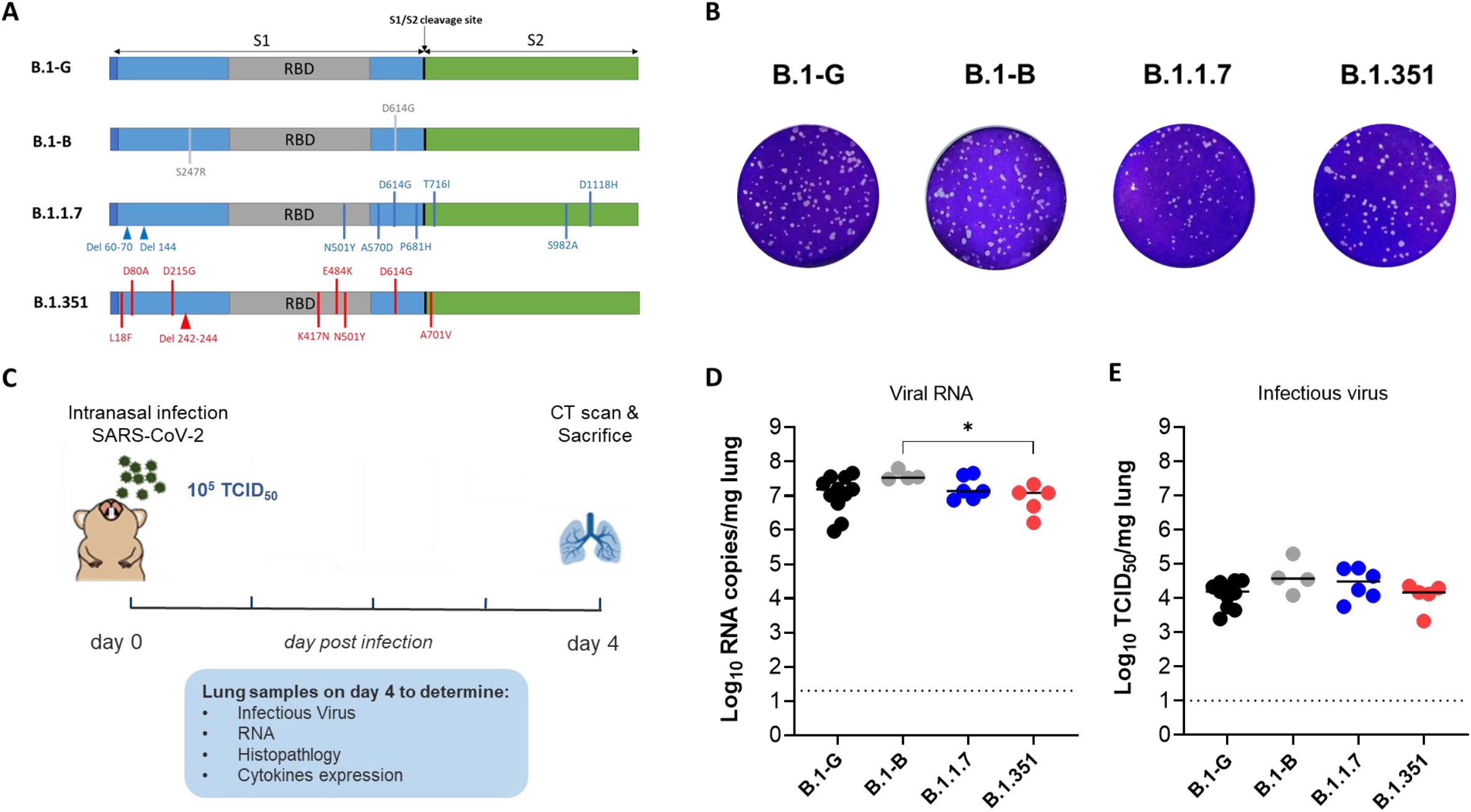
Characterization of the *in vitro* and *in vivo* replication of different SARS-CoV-2 variants. (A) Graphical representation for the SARS-CoV-2 spike gene showing the genotypic difference between the B.1-G, B.1-B, B.1.1.7 and B.1.351 SAR-CoV-2 variants. (B) Plaque phenotype of B.1-G, B.1-B, B.1.1.7 and B.1.351 SAR-CoV-2 variants in Vero E6 cells. (C) Set-up of the Syrian hamsters infection study. (D) Viral RNA levels in the lungs of hamsters infected with 10^5^ TCID_50_ of B.1-G (n=11), B.1-B (n=4), B.1.1.7 (n=6) or B.1.351 (n=5) SAR-CoV-2 variants on day 4 post-infection, p.i are expressed as log_10_ SARS-CoV-2 RNA copies per mg lung tissue. Individual data and median values are presented. (E) Infectious viral loads in the lungs of hamsters infected with the different SARS-CoV-2 variants at day 4 pi are expressed as log_10_ TCID_50_ per mg lung tissue. Individual data and median values are presented. Data were analyzed with the Mann−Whitney U test. *P < 0.05.

Syrian hamsters are a relevant small animal model to study the infectivity and pathogenesis of clinical SARS-CoV-2 isolates.^15,18–20^ We show that such VoC (isolated in 2021) efficiently infect the lower respiratory tract of hamsters following intranasal inoculation, resulting in a pathology that is very similar to that observed after infection with SARS-CoV-2 from earlier more basal lineages (originating from early 2020). In line with clinical evidence from patients infected with VoC B.1.1.7 and B.1.351, major differences in disease outcome with infections caused with the early 2020 isolates were not observed. Nonetheless, animals infected with VoC B.1.1.7 presented with a particularly elevated expression of proinflammatory cytokines, yet with no obvious impact on further aggravation of lung pathology. Overall, we demonstrate that hamsters can serve as relevant preclinical model that can be used to assess the infectivity of clinical SARS-CoV-2 isolates including that of current VoC. This model will also serve as an important tool to study the efficacy of current and future vaccines^17,21,22^ and therapies.^16,23–25^

## Methods

### Virus isolation and virus stocks

All virus-related work was conducted in the high-containment BSL3 facilities of the KU Leuven Rega Institute (3CAPS) under licenses AMV 30112018 SBB 219 2018 0892 and AMV 23102017 SBB 219 2017 0589 according to institutional guidelines.

The basal Severe Acute Respiratory Syndrome-related Coronavirus 2 (SARS-CoV-2) strains isolated in early 2020, i.e. strain BetaCov/Belgium/GHB-03021/2020 (EPI_ISL_407976; 2020-02-03)^26^ and strain Germany/BavPat1/2020 (also referred to as BavPat-1, EPI_ISL_406862; 2020-01-28)^27^ for simplicity called B.1-G and B.1-B, respectively, have been described elsewhere. B.1-G is most closely related to the prototypic Wuhan-Hu-1 2019-nCoV (GenBank accession number MN908947.3) strain;^15^ B.1-B contains the secondary D614G Spike mutation (Fig. 1A). B.1-B was a generous gift of Prof. Christian Drosten, Department of Virology, University Hospital Charité, Berlin, Germany.

SARS-CoV-2 strains belonging to the VoC UK and South African lineages B.1.1.7 (hCoV-19/Belgium/rega-12211513/2020; EPI_ISL_791333, 2020-12-21) and B.1.351 (hCoV-19/Belgium/rega-1920/2021; EPI_ISL_896474, 2021-01-11) were each isolated from nasopharyngeal swabs taken from travelers returning to Belgium in December 2020 and January 2021, respectively; B.1.1.7 from a health subject and B.1.351 from a patient with respiratory symptoms. All strains B.1-G, B.1.1.7 and B.1.351 isolated in house were subjected to sequencing on a MinION platform (Oxford Nanopore)^28^ directly from the nasopharyngeal swabs.

All virus stocks were grown on Vero E6 cells and median tissue culture infectious doses (TCID_50_) defined by titration using the Spearman-Kärber method as decribed.^15^ Virus stocks used throughout this study were from early passages (p); B.1-G and B.1-B from p3, and B.1.1.7 and B.1.351 from p2. Respective Spike gene sequences were confirmed by Sanger sequencing of RT-PCR amplicons of (*i*) virus RNA isolated from virus stocks prior to inoculation and (*ii*) virus RNA isolated from infected hamster lungs 4 dpi. All amino acid variants observed in the Spike protein sequences of the virus strains under study are summarized in Fig. 1A.

### Infection of hamsters

Housing and experimental infections of hamsters have been described^15–17^ and conducted under supervision of the ethical committee of KU Leuven (license P015-2020). In brief, 6 to 8 weeks old female Syrian hamster (*Mesocricetus auratus*) were sourced from Janvier Laboratories and kept per two in individually ventilated isolator cages. Animals were anesthetized with ketamine/xylazine/atropine and intranasally infected with 50 µL of virus stock containing approximately 1 × 10^5^ TCID_50_ (25 µL in each nostril) and euthanized 4 days post infection (dpi) for sampling of the lungs and further analysis.

### Virological and cytokine expression analysis

Virus loads were determined by titration and RT-qPCR from lung homogenates exactly as described in detail before.^15–17^ Analysis of differential expression of hamster interleukin (IL)-6, IL-10, interferon (IFN)-γ, INF-λ, IP-10, TNF-α, MX-2 and ACE by quantitative RT-qPCR has been described.^17^

### Micro-CT and Image Analysis

Micro-CT data for the lungs of free-breathing hamsters were acquired on a Skyscan 1278 system (Bruker Belgium) and analyzed as described before.^15–17^ In brief, hamsters were scanned in supine position under isoflurane anesthesia producing expiratory weighted three-dimensional data sets with 50-μm isotropic reconstructed voxel size^29^ in an approximately 3 min scanning time.

Visualization and quantification of reconstructed micro-CT data were performed with as primary outcome measure a semiquantitative scoring of micro-CT data.^29^ Visual observations were blindly scored on five different, predefined transversal tomographic sections for both lung and airway disease by two independent observers, averaged, and scores for the five sections summed up to obtain a score from 0 to 10, reflecting severity of lung and airway abnormalities compared to healthy control hamsters. As secondary measure, imaging-derived biomarkers (nonaerated, aerated and total lung volume) were quantified as before.^15–17,30^

### Pathology assessment by histology

Assessment of lung pathology was performed as before.^15–17^ In brief, for histological examination, the lungs were fixed overnight in 4% formaldehyde, embedded in paraffin and tissue sections (5 μm) after staining with H&E scored blindly for lung damage (cumulative score of 1 to 3 each for congestion, intra-alveolar hemorrhage, apoptotic bodies in bronchial epithelium, necrotizing bronchiolitis, perivascular edema, bronchopneumonia, perivascular inflammation, peribronchial inflammation, and vasculitis).

## Statistical analysis

All statistical analyses were performed using GraphPad Prism 9 software (GraphPad, San Diego, CA, USA). Results are presented as means ± SEM or medians ± IQR as indicated. Statistical differences between groups were analyzed using Kruskal-Wallis with Dunn’s multiple comparisons test, or two-tailed Mann-Whitney U test for pairwise comparison, and considered statistically significant at p-values ≤0.05 (* p≤0.05, ** p≤0.01, *** p≤0.001).

## Data Statement

All of the data generated or analyzed during this study are included in this published article.

## Results

To investigate the infectivity and pathogenesis of human-adapted VoC B.1.1.7 and B.1.351 (Fig. 1A,B) in hamsters, 6-8 weeks old female Syrian hamsters^15^ were intranasally infected with 50 µL containing approximately 1 × 10^5^ TCID_50_ of either basal lineages (B.1-G, B.1-B) or VoC (B.1.1.7, B.1.351) SARS-CoV-2 (Fig. 1C). Prototypic strains B.1-G^15^ and B.1-B^27^ from early 2020 were included as comparators, the latter containing a spike D614G substitution found in early European lineages and linked to more efficient transmission.^31,32^ At day four post-infection (4 dpi), SARS-CoV-2 replication (Fig. 1D, E), pathology (Fig. 2 and 3), and cytokine expression levels were determined in the lung tissue (Fig. 4 and S2).

**Fig 2.**
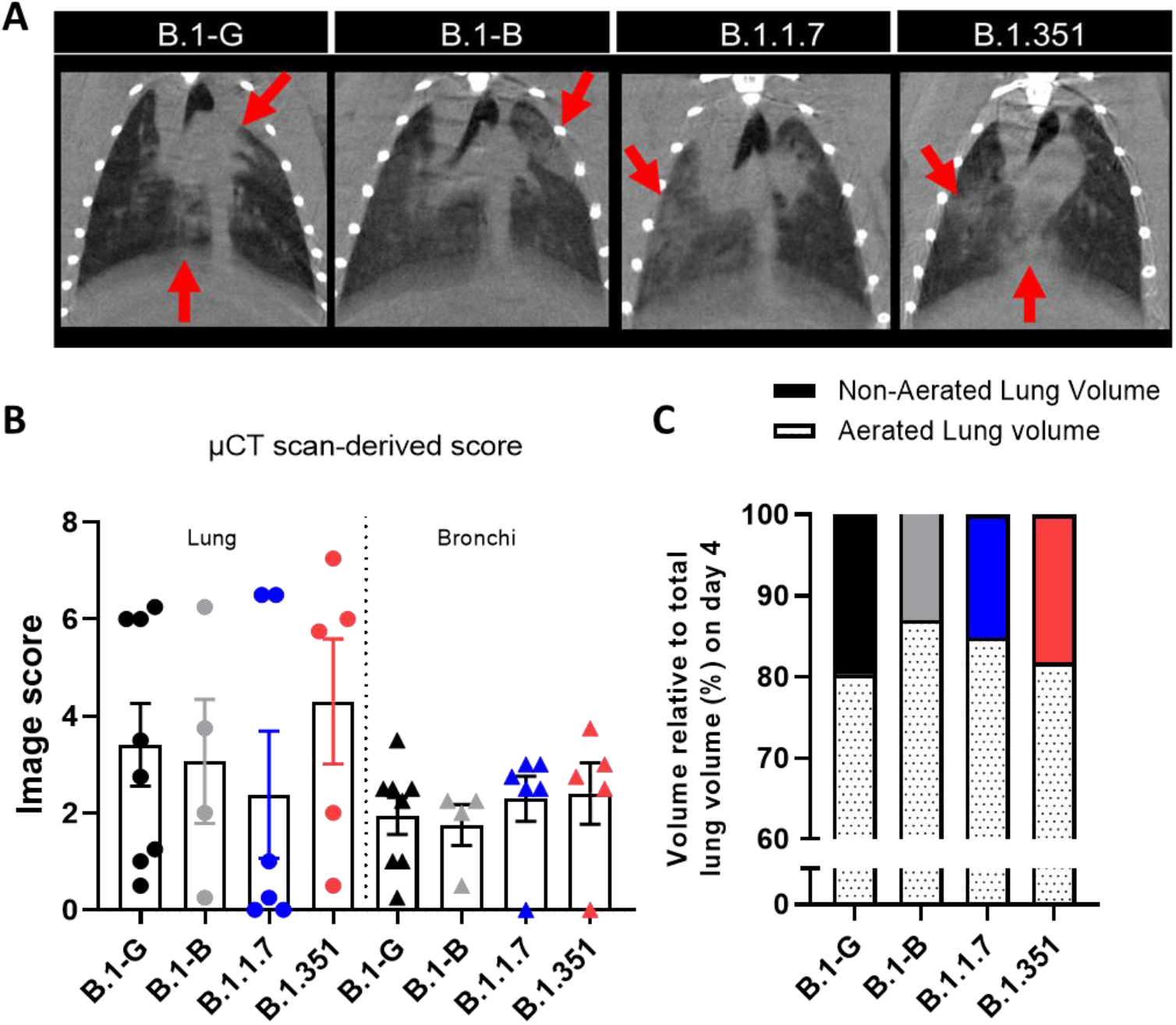
Micro-CT analysis of lung disease burden in Syrian hamsters infected with different SARS-COV-2 variants. (A) Representative coronal lung micro-CT images of hamsters infected with B.1-G, B.1-B, B.1.1.7 or B.1.351 SARS-CoV-2 variants at day 4 pi. Red arrows indicate examples of pulmonary infiltrates. (B) Quantification of the micro-CT-derived lung and bronchi disease scores in hamsters infected with 10^5^ TCID_50_ of B.1-G (n=8), B.1-B (n=4), B.1.1.7 (n=6) or B.1.351 (n=5) SAR-CoV-2 variants on day 4 post-infection. Individual data per hamster are shown and bars represent means ± SEM. (C) Micro-CT-derived non-aerated lung volume (reflecting the tissue lesion volume) and aerated lung volume relative to total lung volume of hamsters infected with the different SARS-CoV-2 variants.

**Fig 3.**
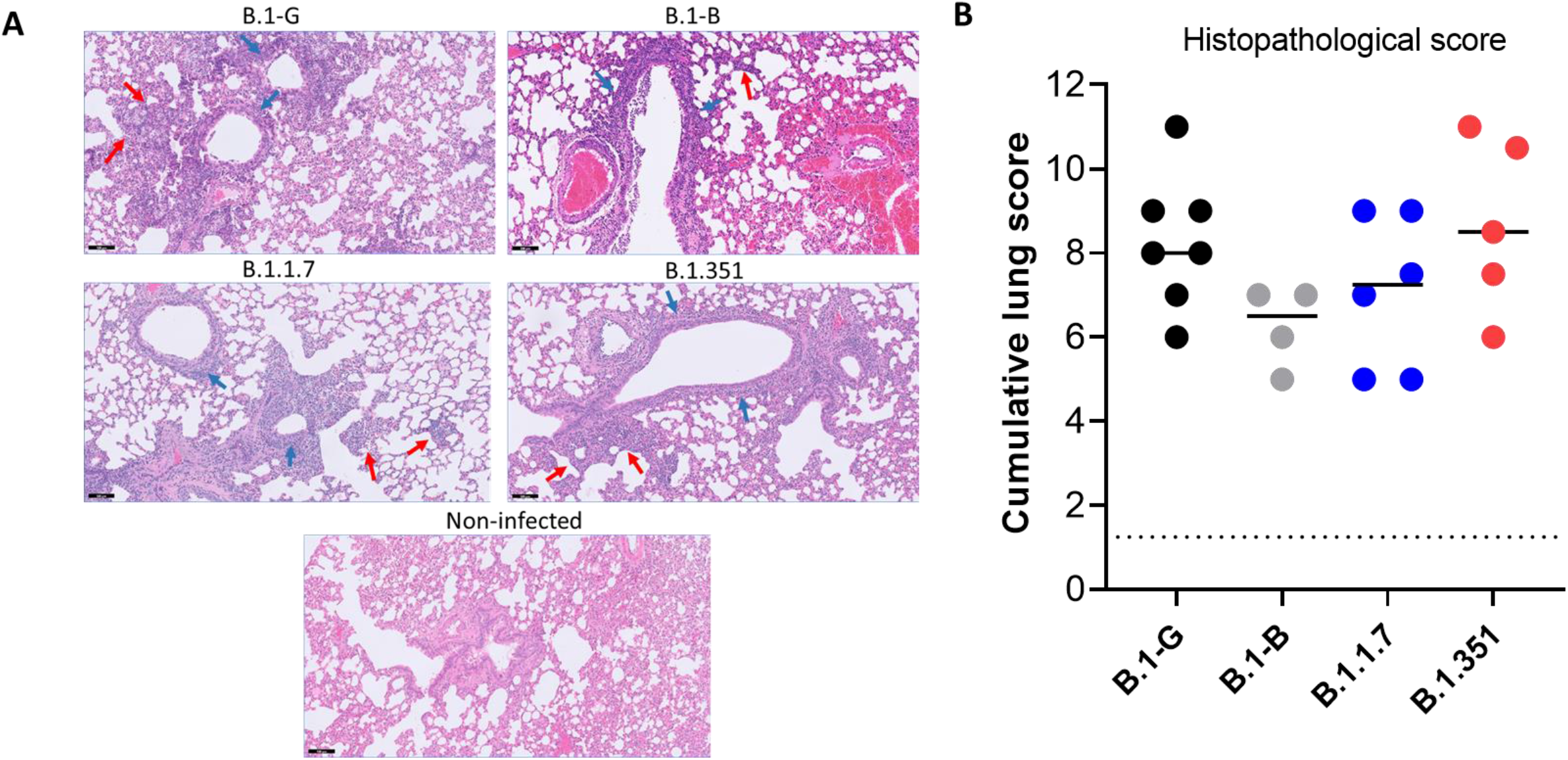
Histopathology of lungs of Syrian hamsters infected with different SARS-CoV-2 variants. (A) Representative H&E images of lungs of hamsters infected with B.1-G, B.1-B, B.1.1.7 or B.1.351 SARS-CoV-2 variants at day 4 pi. The lungs of hamsters infected with any of the SARS-CoV-2 variants show peri-bronchial inflammation (blue arrows) and bronchopneumonia in the surrounding alveoli (red arrows), whereas the lungs of non-infected hamster appear normal. Scale bars, 100 μm. (B) Cumulative severity score from H&E stained slides of lungs from hamsters infected with the different SARS-CoV-2 variants. Individual data and median values are presented and the dotted line represents the median score of untreated non-infected hamsters.

**Fig 4.**
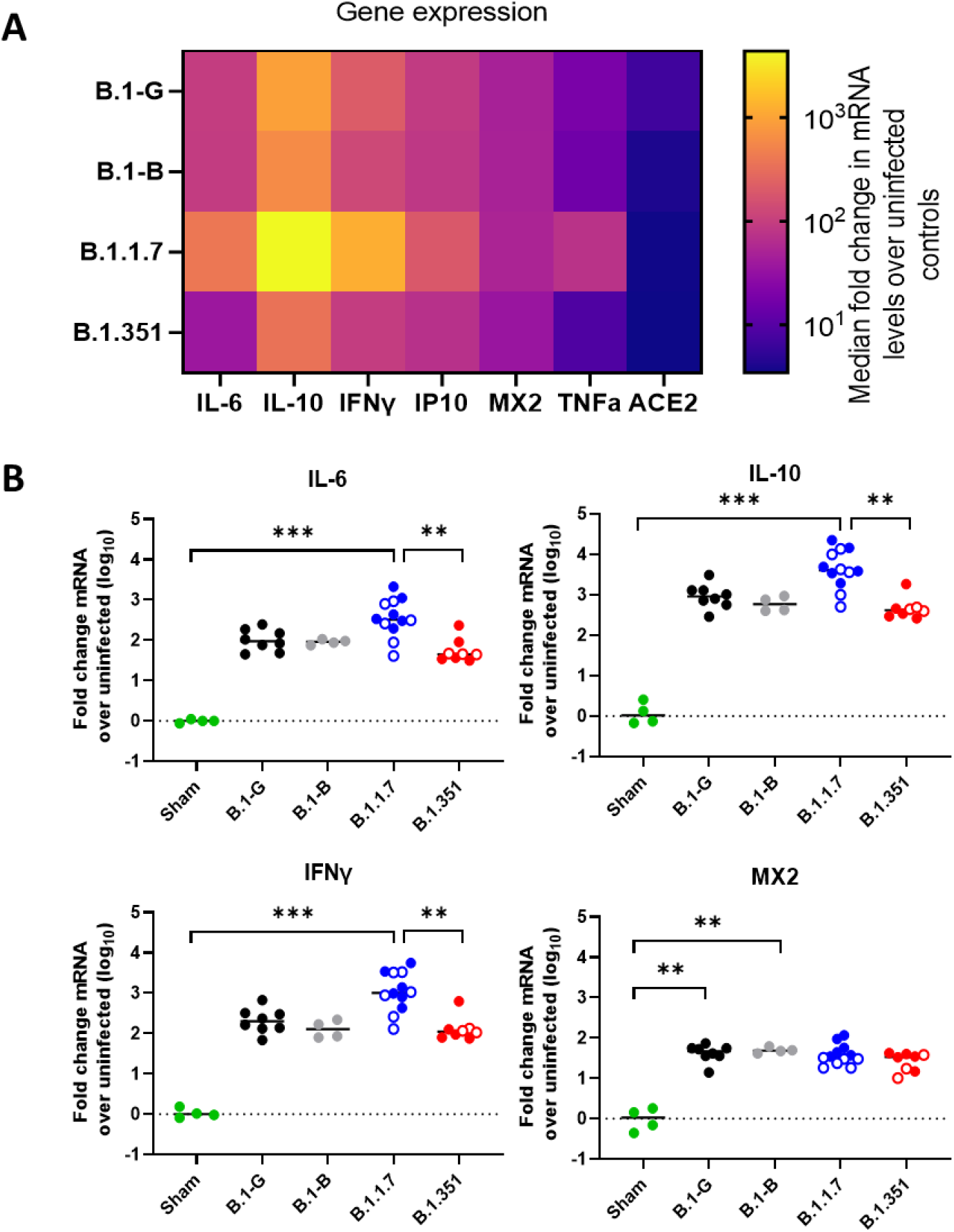
Expression profiles of selected antiviral, pro-inflammatory, and cytokine genes in the lungs after infection with different SAR-CoV-2 variants. (A) Heat map showing differential expression of selected antiviral, pro-inflammatory, and cytokine genes in the lungs after infection with the different SARS-CoV-2 variants relative to non-infected control hamsters. (B) RNA levels for IL-6, IL-10, IFN-γ and MX2 were determined by RT-qPCR on lung extracts from hamsters infected with B.1-G (n=8), B.1-B (n=4), B.1.1.7 (n=12) or B.1.351 (n=8) SAR-CoV-2 variants on day 4 post-infection, normalized for β-actin mRNA levels, and fold changes over the median of uninfected controls were calculated using the 2(−ΔΔCq) method. Data presented as fold change over non-infected control. Closed circles represent hamsters infected with SARS-CoV2 inoculum of 10^5^ TCID50 for all variants, whereas open circles represent 10^4^ TCID_50_ inoculum and 10^2^ TCID_50_ inoculum for B.1.1.7 and B.1.351 variants, respectively. Statistical significance between variants was calculated by Kruskal–Wallis with two-sided Dunn’s post hoc test. **P < 0.01, ***P < 0.001.

Viral RNA loads in the range of 10^7^ RNA copies/mg of lung tissue were consistently detected at 4 dpi for all four strains, with a modest though significant ∼0·5log_10_ decrease in the group infected with B1.351 as compared to B.1-B (Fig. 1D). Infectious virus titers were around 10^4^ TCID_50_/mg of lung tissue for all four strains, with no significant differences between them (Fig. 1E); including for the prototypic 2020 strains, i.e. the D614G containing B.1-B strain and the most basal B.1-G strain, as reported before.^31^ Likewise, when B.1.1.7 and B.1.351 were administered at a lower dose of 10^4^ TCID_50_ and 10^2^ TCID_50_, respectively, similar levels of viral RNA loads and infectious virus titers were detected as with the inoculum of 10^5^ TCID_50_ (data not shown), in line with the high susceptibility of hamsters for clinical SARS-CoV-2 isolates.^20,33^ As documented before for the B.1-G strain in young female hamsters,^15,16^ no overt signs of disease or distress were observed throughout the study, and in general, none of the hamsters showed a marked weight loss (i.e. more than 7%) in any of the infected groups (Fig. S1). Whether or not of biological relevance, some differences in weight evolution could be calculated in a pairwise comparison of groups B.1-B (2% decrease between day 0 and 4 pi) and B.1.1.7 (3% increase) (Fig. S1). Taken together, these results clearly show that VoC B.1.1.7 and B1.351 are able to replicate efficiently in the lower respiratory tract in Syrian hamsters and this to a comparable extent as strains of basal lineages.

Basal lineages and VoC SARS-CoV-2-induced lung pathology was assessed by microcomputed tomography (micro-CT) at 4 dpi immediately prior to euthanasia (Fig. 2) and by post-mortem histopathological analysis (Fig.3). Both semi-quantitative scores and quantitative biomarkers derived from micro-CT thorax scans show similar lung disease burden (pulmonary infiltrates and bronchial dilation)^16^ between the hamsters infected with the different SARS-COV-2 variants.

Hematoxylin/eosin (H&E)-stained images of lungs of hamsters infected with the four variants revealed similar pathological signs with peri-bronchial inflammation and bronchopneumonia in the surrounding alveoli (Fig. 3A). The cumulative histopathological lung scores were 6- to 8-fold higher than the baseline score in untreated, non-infected hamsters (median score of 1·25), with no significant differences between the strains (Fig. 3B). Taken together, both micro-CT and histopathology reveal that the VoC’s induce lung pathology in the hamsters to a similar extent as the basal lineages do, with the same signs of bronchopneumonia and peribronchial inflammation as previously reported.^15,16^

Additionally, the expression levels of a panel of cytokines linked to COVID-19 in humans^34^ were measured in the lung tissue of infected hamsters (Fig. 4). Infection with either of the four strains resulted in an up-regulation of IL-6, IL-10, IFN-λ, IFN-γ, IP-10, MX-2, and TNF-α expression in the range of 10- to 1000-fold compared to non-infected hamsters (Fig. 4, Fig. S2). Notably, IL-6, IL-10and IFN-γ, but not MX-2 expressions were most pronouncedly up-regulated in the B1.1.7 VoC-infected group as compared to the three other strains, with a significant increase compared to B1.351 VoC (Fig. 4B). Similar to lung viral loads (see above), cytokine expression levels were hardly affected by lowering the inoculated virus dose (Fig. 4B and Fig. S2; open circles), arguing for readily saturated conditions. ACE2 receptor expression levels^35^ remained rather unaffected and similar between all four strains (Fig. S2). If at all, a modest up-regulation appeared to occur in lungs of hamsters infected with the basal B.1-G and B.1-B lineages, yet not with the VoC’s. Thus at least at this level, this does not explain differences in epidemiology, transmission, infectivity or virulence as observed or suspected in humans.

## Discussion

SARS-CoV-2 genetic diversification was initially considered slow as the virus was spreading in the first months around the globe.^36^ However, recently more and more variants are emerging and start dominating regional epidemics in widespread populations.^37^ Notably, mutations found in the SARS-CoV-2 spike protein that may be associated with (*i*) more efficient human-to-human transmission, (*ii*) increase virulence,^38^ and/or (*iii*) escape from naturally acquired^7^ or vaccine-induced^9,10^ immunity raise concerns. The role of such mutations have been experimentally addressed, mostly using synthetic viruses generated by targeted mutagenesis. These studies revealed the link between certain mutations and particular phenotypes such as (i) an enhanced transmission potential associated with D614G,^31^ (ii) an increase in virulence associated with N501Y,^39^ and (iii) an impact of E484K on neutralization by post-vaccination sera.^40,41^ We set out to study the infection and pathogenesis of two prototypic VoC B.1.1.7 and B.1.351 (local isolates from the UK and the South African lineages) in the Syrian hamster model using original low passage clinical isolates.

All clinical SARS-CoV-2 isolates studied here, including the VoC B.1.1.7 and B.1.351, replicated efficiently and consistently to high viral loads in hamster lungs (Fig. 1D, E), causing a pathology that resembles bronchopneumonia in COVID patients. However, despite uniformly high viral replication, some variation in disease severity was observed as scored by micro-CT imaging (Fig. 2), histopathology scoring (Fig. 3), and cytokine expression profiles (Fig. 4 and Fig. S2) both (*i*) between study groups as well as (*ii*) among individual animals within particular study groups. At first glance, the hamster model may lack sensitivity to discriminate differences in the replication fitness of SARS-CoV-2 from different lineages, in particular when using direct intranasal infection with high-titred inocula.^31^ Further subtle differences may have been obscured by a lack of statistical power because of the fairly low number of animals per experimental group (n=4-8), despite well-established precedent.^19,20^ However, we^15^ and others^42,43^ have demonstrated before that strain-specific differences may translate into readily detectable changes in pathology in the hamster model. Of note, the particular virus strains used in former proof-of-principle studies contained tissue culture-adaptive mutations (deletions at the S1/S2 junction close to the furin cleavage site).^15,42,43^ By contrast, when low-passage clinical isolates are used for both the basal and VoC lineages (as is the case in the current study), marked quantitative differences cannot be observed. Of note, direct sequencing of viral RNA isolated from infected lungs did not reveal any further sequence evolution in the spike gene during the 4 day course of infection; if at all, a purifying selection and loss of S1/S2 junction variants present in the B.1.351 inoculum.

Nevertheless, our overall observations are fully in line with and hence replicate two key findings from human clinical experience. Firstly, despite some ongoing debate,^38^ there seem to be no obvious differences in pathogenicity associated with recently emerging VoC’s *versus* earlier basal SARS-CoV-2 lineages; definitely, no convergent evolution towards a markedly higher virulence approaching or comparable to what is the case for the highly pathogenic SARS-CoV-1 and MERS coronaviruses.^44^ Secondly, also in humans, SARS-CoV-2 infection shows a high variability in disease severity, clinical presentations and outcome in individual patients suffering from COVID-19.^34^ Some of this variability observed in humans may be mirrored in the range and quantitative fluctuation of disease parameters as measured in the hamster model (Fig. 2-4, Fig. S1 and S2). In our study we focused on a single time point for analysis (4 dpi), previously established to represent a peak of B.1-G virus replication.^15,16^ Obviously, also others reported significant time-, dose-dependent and inter-individual variation in virus kinetics and pathology.^20,31,33^ In conclusion, as far as can be judged from our thorough multi-parameter analysis, VoC B.1.1.7 and B.1.351 compare well in their infectivity and pathogenicity to earlier basal SARS-CoV-2 isolates.

Despite growing insight, it is not known what drives the evolution of SARS-CoV-2. The emergence of future VoC may be driven by any of the following factors: by random selection (founder effects), fitness at the population level (favoring transmission), or viral escape under host immune pressure (antigenic drift), or development of drug resistance under future antiviral therapy. Whatever the cause, as a consequence, any upcoming VoC may spark future COVID-19 epidemics. In an urgent need to characterize current VoC and in anticipation of future needs, the robust hamster model described here will allow to preclinically asses (*i*) virus transmission, (*ii*) vaccine efficacy, and (*iii*) evaluation of pharmacological interventions that target B.1.1.7 and B.1.351 as well as expected future VoC. Importantly, our primary analysis of infection with B1.1.7 (UK) and B.1.351 (South African) SARS-CoV-2 VoC in hamsters does not reveal evidence for a largely altered phenotype confined to these lineages.

## Acknowledgements

We thank Carolien De Keyzer, Lindsey Bervoets, Thibault Francken, Birgit Voeten, Dagmar Buyst, Niels Cremers for excellent technical assistance with animal experimentation and molecular sample analysis. We also thanks Bo Corbeels and Kathleen Van den Eynde for histopathology samples processing and staining. SARS-CoV-2 strain BavPat-1 was a generous gift of Prof. Christian Drosten, Department of Virology, University Hospital Charité, Berlin, Germany.

The authors acknowledge funding by the Flemish Research Foundation (FWO) emergency Covid-19 fund (G0G4820N) and the FWO Excellence of Science (EOS) program (No. 30981113; VirEOS project), the European Union’s Horizon 2020 research and innovation program (No 101003627; SCORE project and No 733176; RABYD-VAX consortium), the Bill and Melinda Gates Foundation (INV-00636), KU Leuven Internal Funds (C24/17/061) and the KU Leuven/UZ Leuven Covid-19 Fund (COVAX-PREC project). L.S. acknowledges support by an aspirant mandate from the FWO (1186121N). X.Z. received funding of the China Scholarship Council (grant No.201906170033). K.D. acknowledges grant support from KU Leuven Internal Funds (C3/19/057 Lab of Excellence).

## Author Contributions

R.A., C.S.F., S.J.F.K., R.B., L.D., K.D. and J.N. designed the studies; R.A., R.B., L.S.F., S.J.F.K., X.Z. and L.S. performed the experiments; R.A., R.B., L.S., C.S.F., L.S.F, G.V.V. and B.W. analyzed data; R.A., R.B. and L.S.F. designed the figures; J.N. and K.D. provided advice on the interpretation of data; K.D., C.S.F., R.B. and R.A. wrote the paper with input from co-authors; P.M. isolated and initially characterized variants; R.A., L.D., S.J.F.K., G.V.V., K.D. and J.N. supervised the study; J.N. and K.D. acquired funding.

## Figure Legends

**Fig S1.**
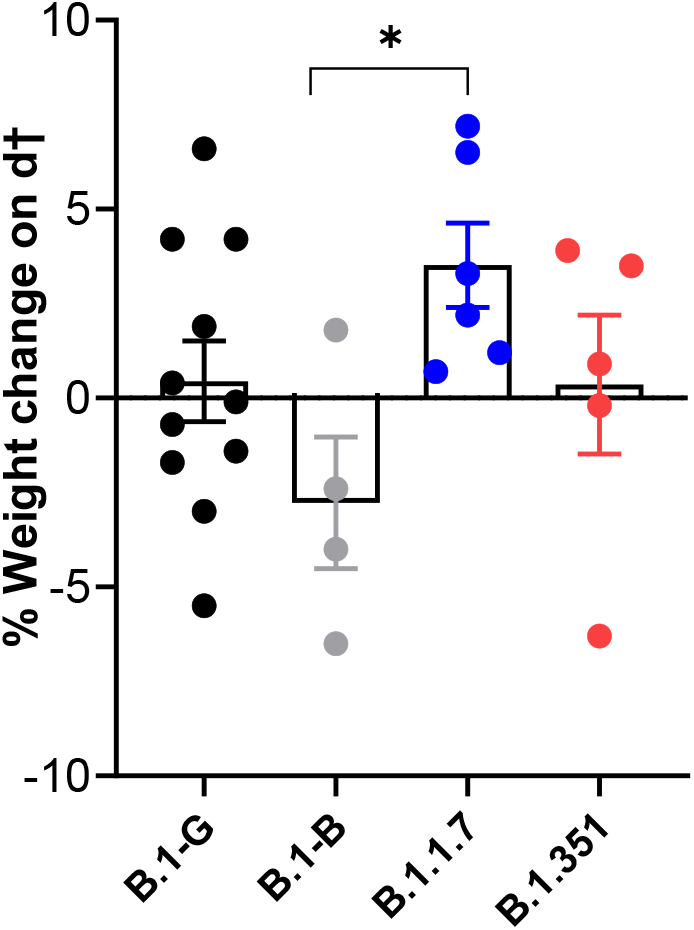
Weight change in hamsters infected with different SARS-CoV-2 variants. Weight change at day 4 post-infection with 10^5^ TCID_50_ of B.1-G (n=11), B.1-B (n=4), B.1.1.7 (n=6) or B.1.351 (n=5) SAR-CoV-2 variants represented in percentage and normalized to the body weight at the time of infection. Bars represent means ± SEM. Data were analyzed with the Mann−Whitney U test. *P < 0.05.

**Figure S2.**
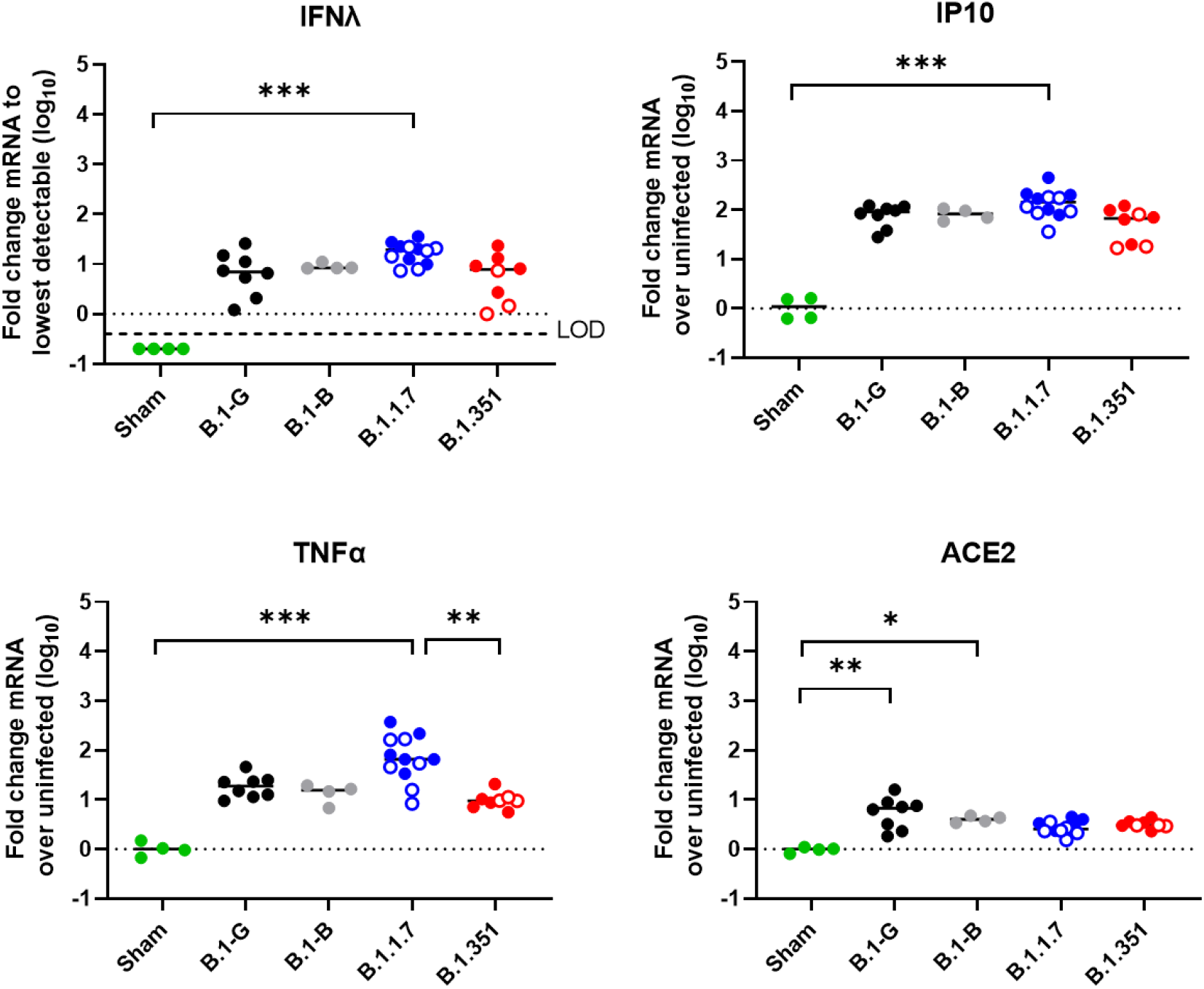
Expression profiles of selected antiviral, pro-inflammatory, and cytokine genes in the lungs after infection with different SAR-CoV-2 variants. RNA levels for each marker gene were determined by RT-qPCR on lung extracts from hamsters infected with B.1-G (n=8), B.1-B (n=4), B.1.1.7 (n=12) or B.1.351 (n=8) SAR-CoV-2 variants on day 4 post-infection, normalized for β-actin mRNA levels, and fold changes over the median of uninfected controls were calculated using the 2(−ΔΔCq) method. Only for IFN-λ, where all uninfected control animals had undetectable RNA levels, fold changes were calculated over the lowest detectable value. Data presented as fold change over non-infected control. Closed circles represent hamsters infected with SARS-CoV2 inoculum of 10^5^ TCID_50_ for all variants, whereas open circles represent 10^4^ TCID_50_ inoculum and 10^2^ TCID_50_ inoculum for B.1.1.7 and B.1.351 variants, respectively. Statistical significance between variants was calculated by Kruskal– Wallis with two-sided Dunn’s post hoc test. *P < 0.05, **P < 0.01, ***P < 0.001.

## References

1 O’Toole Á, Hill V, Pybus OG, et al. Tracking the international spread of SARS-CoV-2 lineages B.1.1.7 and B.1.351/501Y-V2. Virological. 2021; Published online January 14. https://virological.org/t/tracking-the-international-spread-of-sars-cov-2-lineages-b-1-1-7-and-b-1-351-501y-v2/592 (accessed Feb 24, 2021).

2 O’Toole Á, Hill V. Lineage B.1.1.7. 2021; Published online February 25. https://cov-lineages.org/global_report_B.1.1.7.html (accessed Feb 26, 2021).

3 Rambaut A, Holmes EC, O’Toole Á, et al. A dynamic nomenclature proposal for SARS-CoV-2 lineages to assist genomic epidemiology. Nat Microbiol 2020; 5: 1403–7.

4 O’Toole Á, Hill V. Lineage B.1.351. 2021; Published online February 25. https://cov-lineages.org/global_report_B.1.351.html (accessed Feb 26, 2021).

5 Tegally H, Wilkinson E, Giovanetti M, et al. Emergence and rapid spread of a new severe acute respiratory syndrome-related coronavirus 2 (SARS-CoV-2) lineage with multiple spike mutations in South Africa. medRxiv 2020. 2020.12.21.20248640.

6 O’Toole Á, Hill V. Lineage P.1. 2021; Published online February 25. https://cov-lineages.org/global_report_P.1.html (accessed Feb 26, 2021).

7 Sabino EC, Buss LF, Carvalho MPS, et al. Resurgence of COVID-19 in Manaus, Brazil, despite high seroprevalence. Lancet 2021; 397: 452–5.

8 Luan B, Wang H, Huynh T. Molecular Mechanism of the N501Y Mutation for Enhanced Binding between SARS-CoV-2’s Spike Protein and Human ACE2 Receptor. bioRxiv 2021. 2021.01.04.425316.

9 Mahase E. Covid-19: South Africa pauses use of Oxford vaccine after study casts doubt on efficacy against variant. BMJ 2021; 372: 372.

10 Mahase E. Covid-19: Novavax vaccine efficacy is 86% against UK variant and 60% against South African variant. BMJ. 2021; 372: 296.

11 Greaney AJ, Starr TN, Gilchuk P, et al. Complete Mapping of Mutations to the SARS-CoV-2 Spike Receptor-Binding Domain that Escape Antibody Recognition. Cell Host Microbe 2021; 29: 44-57.e9.

12 Wang Z, Schmidt F, Weisblum Y, et al. mRNA vaccine-elicited antibodies to SARS-CoV-2 and circulating variants. Nature 2021. DOI:10.1038/s41586-021-03324-6.

13 Wise J. Covid-19: The E484K mutation and the risks it poses. BMJ 2021; 372: 359.

14 Hee Ko S, Bayat Mokhtari E, Mudvari P, et al. High-Throughput, Single-Copy Sequencing Reveals SARS-CoV-2 Spike Variants Coincident with Mounting Humoral Immunity during Acute COVID-19. bioRxiv 2021. 2021.02.21.432184.

15 Boudewijns R, Thibaut HJ, Kaptein SJF, et al. STAT2 signaling restricts viral dissemination but drives severe pneumonia in SARS-CoV-2 infected hamsters. Nat Commun 2020; 11: 1–10.

16 Kaptein SJF, Jacobs S, Langendries L, et al. Favipiravir at high doses has potent antiviral activity in SARS-CoV-2−infected hamsters, whereas hydroxychloroquine lacks activity. Proc Natl Acad Sci USA 2020; 117: 26955–65.

17 Sanchez-Felipe L, Vercruysse T, Sharma S, et al. A single-dose live-attenuated YF17D-vectored SARS-CoV-2 vaccine candidate. Nature 2020; 590: 320–5.

18 Muñoz-Fontela C, Dowling WE, Funnell SGP, et al. Animal models for COVID-19. Nature 2020; 586: 509–15.

19 Chan JF-W, Zhang AJ, Yuan S, et al. Simulation of the clinical and pathological manifestations of Coronavirus Disease 2019 (COVID-19) in golden Syrian hamster model: implications for disease pathogenesis and transmissibility. Clin Infect Dis 2020; 71: 2428–2446.

20 Imai M, Iwatsuki-Horimoto K, Hatta M, et al. Syrian hamsters as a small animal model for SARS-CoV-2 infection and countermeasure development. Proc Natl Acad Sci USA 2020; 117: 16587–95.

21 Krammer F. SARS-CoV-2 vaccines in development. Nature 2020; 586: 516–27.

22 Tostanoski LH, Wegmann F, Martinot AJ, et al. Ad26 vaccine protects against SARS-CoV-2 severe clinical disease in hamsters. Nat Med 2020; 26: 1694–700.

23 Abdelnabi R, Foo CS, Kaptein SJF, et al. Molnupiravir (EIDD-2801) inhibits SARS-CoV-2 replication in Syrian hamsters model. bioRxiv 2020. 2020.12.10.419242.

24 Vandyck K, Abdelnabi R, Gupta K, et al. ALG-097111, a potent and selective SARS-CoV-2 3-chymotrypsin-like cysteine protease inhibitor exhibits in vivo efficacy in a Syrian Hamster model. bioRxiv 2021. 2021.02.14.431129.

25 Foo CS, Abdelnabi R, Kaptein SJF, et al. Nelfinavir markedly improves lung pathology in SARS-CoV-2-infected Syrian hamsters despite a lack of an antiviral effect. bioRxiv 2021. 2021.02.01.429108.

26 Spiteri G, Fielding J, Diercke M, et al. First cases of coronavirus disease 2019 (COVID-19) in the WHO European Region, 24 January to 21 February 2020. Eurosurveillance 2020; 25: 2000178.

27 Rothe C, Schunk M, Sothmann P, et al. Transmission of 2019-nCoV Infection from an Asymptomatic Contact in Germany. N Engl J Med 2020; 382: 970–1.

28 Vrancken B, Wawina-Bokalanga T, Vanmechelen B, et al. Accounting for population structure reveals ambiguity in the Zaire Ebolavirus reservoir dynamics. PLoS Negl Trop Dis 2020; 14: e0008117.

29 Berghen N, Dekoster K, Marien E, et al. Radiosafe micro-computed tomography for longitudinal evaluation of murine disease models. Sci Rep 2019; 9: 1–10.

30 Vande Velde G, Poelmans J, De Langhe E, et al. Longitudinal micro-CT provides biomarkers of lung disease that can be used to assess the effect of therapy in preclinical mouse models, and reveal compensatory changes in lung volume. DMM Dis Model Mech 2016; 9: 91–8.

31 Hou YJ, Chiba S, Halfmann P, et al. SARS-CoV-2 D614G variant exhibits efficient replication ex vivo and transmission in vivo. Science 2021; 370: 1464–8.

32 Volz E, Hill V, McCrone JT, et al. Evaluating the Effects of SARS-CoV-2 Spike Mutation D614G on Transmissibility and Pathogenicity. Cell 2021; 184: 64-75.e11.

33 Rosenke K, Meade-White K, Letko M, et al. Defining the Syrian hamster as a highly susceptible preclinical model for SARS-CoV-2 infection. Emerg Microbes Infect 2020; 9: 2673–84.

34 Wiersinga WJ, Rhodes A, Cheng AC, et al. Pathophysiology, Transmission, Diagnosis, and Treatment of Coronavirus Disease 2019 (COVID-19): A Review. J. Am. Med. Assoc. 2020; 324: 782–93.

35 Onabajo OO, Banday AR, Stanifer ML, et al. Interferons and viruses induce a novel truncated ACE2 isoform and not the full-length SARS-CoV-2 receptor. Nat Genet 2020; 52: 1283–93.

36 Dearlove B, Lewitus E, Bai H, et al. A SARS-CoV-2 vaccine candidate would likely match all currently circulating variants. Proc Natl Acad Sci USA 2020; 117: 23652–62.

37 Ecdc. Risk related to the spread of new SARS-CoV-2 variants of concern in the EU/EEA-first update. 2021; Published online January 21. https://beta.microreact.org/project/r8vBmatkC9mcfrJJ6bUtNr-cog-uk-2021-01-09-sars-cov-2-in-the-uk/ (accessed Feb 24, 2021).

38 Davies NG, Barnard RC, Jarvis CI, et al. Estimated transmissibility and severity of novel SARS-CoV-2 Variant of Concern 202012/01 in England. medRxiv 2020. 2020.12.24.20248822.

39 Rathnasinghe R, Jangra S, Cupic A, et al. The N501Y mutation in SARS-CoV-2 spike leads to morbidity in obese and aged mice and is neutralized by convalescent and post-vaccination human sera. medRxiv 2021. 2021.01.19.21249592

40 Xie X, Liu Y, Liu J, et al. Neutralization of SARS-CoV-2 spike 69/70 deletion, E484K and N501Y variants by BNT162b2 vaccine-elicited sera. Nat Med 2021. DOI:10.1038/s41591-021-01270-4.

41 Wang P, Liu L, Iketani S, et al. Increased Resistance of SARS-CoV-2 Variants B.1.351 and B.1.1.7 to Antibody Neutralization. bioRxiv 2021. 2021.01.25.428137.

42 Johnson BA, Xie X, Bailey AL, et al. Loss of furin cleavage site attenuates SARS-CoV-2 pathogenesis. Nature 2021. DOI:10.1038/s41586-021-03237-4.

43 Lau S-Y, Wang P, Mok BW-Y, et al. Attenuated SARS-CoV-2 variants with deletions at the S1/S2 junction. Emerg Microbes Infect 2020; 9: 837–42.

44 Zhu Z, Lian X, Su X, et al. From SARS and MERS to COVID-19: A brief summary and comparison of severe acute respiratory infections caused by three highly pathogenic human coronaviruses. Respir. Res. 2020; 21: 1–14

